# Neural basis of cognitive control signals in anterior cingulate cortex during delay discounting

**DOI:** 10.1101/2024.06.07.597894

**Authors:** Jeremy K. Seamans, Eldon Emberly, Shelby White, Mitchell Morningstar, David Linsenbardt, Baofeng Ma, Cristine L. Czachowski, Christopher C. Lapish

## Abstract

Cognitive control involves allocating cognitive effort according to internal needs and task demands. The anterior cingulate cortex (ACC) is hypothesized to play a central role in this process. We investigated the neural basis of cognitive control in the ACC of rats performing an adjusting-amount delay discounting task, with a 4s or 8s delay between lever choice and reward. A reinforcement learning model indicated that decision making on this task can be guided by either a value tracking strategy, requiring a ‘resource-based’ form of cognitive control or a delay-lever biased strategy requiring a ‘resistance-based’ form of cognitive control. This was then tested in vivo by multiple single unit recordings and local field potentials acquired from male rats performing the task. On this task, the behavioral manifestation of resistance-based control is an excessive focus on delayed lever choices which was observed during a substantial portion of 4s but not 8s delay sessions. On a neural level, this was associated with an increase in Theta (6-12Hz) oscillations prior to delay choices which was present exclusively on 4s delay sessions. By contrast, evidence of a resource-based control signal was found in spike trains that closely tracked lever value prior to choice, and was far more prevalent on 8s delay sessions. These data provide candidate neural signatures of ‘resource-based’ versus ‘resistance-based’ forms of cognitive control. While mediated by distinct neural mechanisms, either form could be engaged by individual subjects under different task conditions.

## Introduction

Cognitive effort is commonly thought of as being equivalent to mental exertion, however multiple types of cognitive effort have been identified and formalized^1,2^. A *resourced-based* form of cognitive effort is relevant whenever a task relies on a valuable but depleting resource, such as attention or working memory^1,3,4^. There is also a *resistance-based* form of cognitive effort that is used to overcome some type of internal ‘resistive force’, such as unpleasantness or impatience^2,5^. Both types of cognitive effort are costly and are typically avoided when possible. However, the perceived costs of deploying cognitive effort must be weighed against the consequences of not exerting effort. Such decisions are influenced by a variety of extrinsic and subjective factors, including the task demands, the individual’s propensity for one type of cognitive effort over another, their level of arousal or fatigue, and the urgency of the need that would be satisfied by exerting the effort^2,5^. For these reasons, cognitive effort cannot be measured in purely objective terms but must be inferred from a subject’s behavior or from physiological measures.

Decision-making tasks involving intertemporal choices have commonly been used to study cognitive effort in both humans^4^ and rats^6,7^. These tasks require subjects to choose between accepting a low-value reward immediately or waiting for a higher valued reward to be delivered after a delay. The adjusting amount delay-discounting task^8,9^ adds an additional layer of complexity, as the payout for the immediate option decreases whenever the immediate option is chosen and increases whenever the delayed option is chosen. In order to both maximize reward and minimize waiting, the subject should generally favor the delayed lever option but exploit the immediate lever option whenever its value (i.e. payout) is high. As a result, these tasks require a combination of the two types of cognitive effort mentioned above: The resistance-based form is needed in order to overcome the unpleasantness of waiting out the delays associated with the delayed lever option, whereas the resourced-based form cognitive effort is necessary when the subject must draw on attentional and mnemonic resources to keep track of the value of the immediate lever option to know when its value is high.

Deciding how and when to deploy cognitive effort is a form of cognitive control that is thought to rely primarily on the anterior cingulate cortex (ACC). Cognitive control processes, mediated by thea, dynamically regulate how much and what type of cognitive effort should be deployed by comparing the expected value of the outcome produced by the effort versus the cost of implementing and maintaining that effort^1,10^. In this regard, the cost of cognitive effort is ‘felt’ at a physiological level through changes in emotion and autonomic tone^2,4,11–13^.

The ACC is well suited to track the cost of cognitive effort due to its extensive bilateral connections with regions involved in regulating autonomic tone^14–16^. There is also considerable support for the idea that the ACC is involved in effort-based decision-making more generally as multiple studies have shown that ACC activity represents the degree of expected effort^17–19^ and that ACC lesions in rodents reduce the propensity to make choices that involve physical effort, even if such efforts result in higher rewards^20–22^. In line with this view, inactivation of the ACC reduces the willingness of rats to expend cognitive effort to obtain larger rewards on a variant of the 5-choice serial reaction time task^23^. In that study, cognitive effort was defined based on the relative demands placed on visuospatial attention, which is an example resourced-based cognitive effort.

The neural mechanisms of cognitive control have been studied at a macroscopic level in humans using EEG^24,25^ and theta oscillations have emerged as a candidate signal. Frontomedial theta oscillations originate in the ACC and potentially synchronize ACC with other brain regions when control is deployed^26,27^. Fronotmedial theta exhibits several features that would be expected of a cognitive control signal, as it increases following negative feedback over time in tasks that require sustained effort, and under conditions when a decision is difficult^28,29^. Further, a role for theta oscillations in allocating physical effort has been identified^30,31^. However, the cellular mechanisms within the ACC that control and deploy types of cognitive effort have not been identified.

We sought to address this knowledge gap by recording from ensembles of ACC neurons during the adjusting amount delay-discounting task. If rats opt to make choices based on the relative values of the two levers, they would need to rely on a resource-based form of cognitive effort in order to keep track of the ever-changing value of the immediate lever (ival). This strategy would be expressed as a switch from focusing on the delay option when ival is low to the immediate option when ival is high. Alternatively, a rat may focus solely on waiting out the delay throughout the session. While this strategy forgoes the need for ival tracking, it does require a strong reliance on a resistance-based form of cognitive effort to endure the delays associated with each delay choice. Neural and behavioral correlates of each strategy were assessed using a delay discounting (DD) task.

## Methods

### Subjects and task

For electrophysiology experiments, 10 male Wistar rats were purchased from Envigo (Indianapolis, IN). Animals were acclimated to vivarium conditions, a 12-h reverse light/dark cycle with lights ON at 7:00 PM, for 3 days prior to handling. Animals were then single-housed with ad lib access to food and water for a week and were at least 70 days of age prior to food restriction and habituation to the task. All procedures were approved by the IUPUI School of Science Institutional Animal Care and Use Committee and were in accordance with the National Institutes of Health Guidelines for the Care and Use of Laboratory Animals.

### Apparatus

Behavioral training was performed in eight standard operant boxes (20.3 cm × 15.9 cm × 21.3 cm; Med Associates, St Albans, VT) inside of sound attenuating chambers (ENV-018M; MED Associates, St. Albans, VT). Each box contained one wall with two stimulus lights, two retractable levers that flanked a pellet hopper, and a tone generator. Cue lights were above each lever. A tone generator (2900 Hz) was above the hopper. A house light was on the opposite wall.

All awake-behaving electrophysiological recordings were performed in one custom-built operant box (21.6 cm x 25.7 cm x 52.0 cm). Dimensions, stimuli (including house and cue lights), and retractable levers were all positioned to replicate the conditions of the operant boxes as closely as possible. The floor bars of the custom-built box were made of painted wooden dowels. All metal components of the box were powder coated. MED-PC IV software (Med Associates, St. Albans, VT) was used to all environmental variables (e.g. lever extensions, presses, lights on/off).

### Behavioral Training

Following a week of single housing, animals were handled daily for a week. Animals were food restricted to 85% of their starting free-feeding weight and maintained under this condition for the duration of the experiment except for immediately before and after surgery. Animals received their daily amount of food following testing.

Animals were habituated on Day 1 to the operant chambers for 30 minutes. Shaping procedures began on Day 2 and details for shaping procedures can be found in^32^. In brief, animals were trained to press each lever for a sucrose pellet and environmental variables were introduced in a staged manner over 6 days. Learning the contingency between pressing the lever and receiving a pellet was demonstrated by a minimum of 30 successful reinforced lever responses for each lever, which was typically completed within 6 days.

On days 7 and 8 training on the task with all environmental stimuli was initiated. Illumination of the house light signaled the start of a trial and remained on for 10 seconds. Once extinguished, both levers extended, and the animal was required to press either lever in order to initiate the start of the trial. No response for 10 seconds resulted in retraction of the levers followed by the illumination of the house light (10 seconds). Once a lever was pressed to initiate the trial, both levers retracted for 1 second. Levers were reinserted into the chamber and lights above each lever were illuminated. A response on either lever was marked with a 100ms tone and a single sucrose pellet delivered, simultaneously. Only the cue light above the chosen lever remained on for the remainder of the trial. The duration of the trials was always 35 seconds. These sessions were terminated either when 30 choices were made or when 35 minutes had elapsed. Over sessions 7 and 8, lever preference bias was determined for each animal.

### Delay Discounting Task

The adjusting reward DD task was a modified version of the procedure^32^ which was originally adapted from Oberlin and Grahame^33^. Stimuli were presented in the exact manner detailed in days 7 and 8 of shaping except that the number of pellets delivered for a given trial was dependent on lever pressing contingencies detailed below. The “delay lever” was assigned to each animal as their non-preferred side. Choosing the delay lever always resulted in the delivery of 6 pellets following some delay (0, 1, 2, 4, or 8s). Choosing the immediate lever resulted in 0-6 pellets delivered immediately (i.e. the adjusting amount lever). The value of the immediate lever (ival), was defined as the number of pellets delivered by the immediate lever. Ival was always set to 3 pellets at the start of each session. On “choice trials” each response on the immediate lever would decrease the number of pellets the immediate lever would dispense on the next trial by one (minimum 0 pellets) whereas a response on the delay lever would increase the number of pellets the immediate lever would dispense on the next trial by one (max 6 pellets). “Forced trials” were implemented for the immediate and delay levers, where two consecutive responses on the same lever would result in a forced trial for the non-chosen lever on the next trial. If an animal did not lever press on the forced trial, the forced trial would be presented again until the lever was pressed. The animal had to eventually make a response on the forced trial to return to choice trials. There was no effect of forced trials on the value of the immediate lever. Further, choosing the immediate lever did not alter the reinforcement rate throughout the session.

Animals then completed 0, 1, and 2-sec delays in ascending order before surgery. After completing a delay, animals had one day off where no testing occurred. Eight to twelve sessions were given at the 0-sec delay and four sessions at the 1 and 2-sec delay. Reward magnitude discrimination was determined at the 0-sec delay in the standard operant chambers using the 45mg sucrose pellets with an exclusion criterion of 70% (4.2 pellets) of the maximum reward value (6 pellets). Magnitude criteria were meant to assess whether animals understood the lever contingencies before moving forward with subsequent delays, specifically, that there was no penalty for pressing the delay lever at a 0-sec delay.

Once the animals recovered from surgery (see below) they were given a 2-sec delay ‘reminder session’ before recording neural activity. Each recording session consisted of 40 choice trials completed within 45mins and used 20mg sucrose pellets.

### Electrophysiology Surgical Preparation & Implantation

Animals were anaesthetized with isoflurane gas (2% at 4L/h) until sedated, at which point they were placed in a stereotaxic frame and maintained on 0.3-0.5% isoflurane for the duration of the surgery. Artificial tears were then applied. Subsequently, fur was shaved and the skin at the incision site was sanitized with three rounds of both 70% EtOH and betadine before applying a local anesthetic (Marcaine; 5mg/kg s.c.). An anti-inflammatory (Ketofen; 5mg/kg dose s.c.) and antibiotic (Cefazolin; 30mg/kg s.c.) were injected at the nape of the neck (anti-inflammatory and antibiotic) before beginning the incision. Once the skull was exposed and cleaned of blood, bregma-lambda coordinates were identified. Prior to implantation of Cambridge Probes, four anchoring screws were inserted.

A small, rectangular craniotomy was performed over the right hemisphere of MFC (AP: 2.8, ML: 0.3 from bregma) followed by a durotomy and cleaning/hydration of the probe insertion site with a sterile saline solution. Additionally, two ground screws were placed above the cerebellum. A Cambridge Neurotech F (n=5), P (n=4), or E-series (n=1) 64-channel silicon probe on a movable drive (Cambridge Neurotech, Cambridge, UK) was lowered to the target site. Mobility of the movable drive was maintained with a coating of antibiotic ointment. Following insertion of Cambridge Probes, a two-compound dental cement was used to adhere implants to anchoring screws. Following completion of surgical procedures, animals were maintained in a clean heated cage and monitored for recovery before being returned to the vivarium. Histological assessments of electrodes placements are provided in ^34^.

### Electrophysiology Equipment

An electrical interface board connected silicon electrodes to an Intan Omnetics headstage (Intan – CA). An Intan RHD SPI cable (Intan – CA) connected the headstage to a Doric Commutator (Doric Lenses – Canada) positioned above the operant apparatus. An OpenEphys (OpenEphys – MA) acquisition system was used to collect all electrophysiological data. Data was streamed from the OpenEphys acquisition box to a compatible desktop computer via a USB 2.0 connection and sampled at 30 kHz. AnyMaze (ANY-maze Behavioral tracking software – UK) was used to collect all behavioral and locomotor data. ANY-maze locomotor data was synchronized with OpenEphys via an ANY-maze AMI connected to an OpenEphys ADC I/O board. Med PC behavioral events were also synchronized to the electrophysiological recordings via an OpenEphys ADC I/O board. Following sessions with diminished signal, electrodes were lowered 50µm following completion of that session in order to allow any drifting of the probe to occur before the next day’s session. The unsupervised portion of spike sorting was performed Kilosort 2.0^35^, followed by supervised cluster evaluation in Phy (https://github.com/cortex-lab/phy). Spike train autocorrelations were evaluated to ensure that minimal refractory period violations were observed and waveforms were consistent with that of an action potential.

### Behavior Clustering

The behavior of each animal on a given session could be summarized by its sequence of lever presses on the free choices. We assigned trials where the delay lever was pressed as +1 and the immediate lever pressed as -1. For an animal on a given session, we took the cumulative sum of its choices as its behavioral profile. We then used hierarchical clustering (using Ward’s method) on the profiles for the sessions with a given delay to identify groups of rats that had similar trends in their choice of levers. This clustering led to identifying 2 groups of different behavior at each delay.

### Stochastic choice and Reinforcement Learning Models

Two types of simulations of the DD task explored how choices were influenced by varying parameters representing an initial lever bias and resource tracking. To explore the impact of a lever bias, stochastic simulations were performed where choices are assigned simply based on probabilities. In this approach, a subject’s delay lever bias was characterized by the parameter, *p*_*D*_, which sets the default probability of picking the delay lever. Levers were picked at random using *p*_*D*_, (i.e. a uniform random number was drawn and it if was less than *p*_*D*_, then the delay lever was chosen, otherwise immediate) and the simulation updated the number of pellets rewarded by the immediate lever according to the task rules. After 40 free choices were achieved, the total number of pellets received on the free choices was reported.

For the case where a subject tracks resources, we assume that they assign a certain reward value to the number of pellets received, which depends both on how they value a single pellet as well as one lever versus the other. To accommodate this a Reinforcement Learning approach was implemented in the model. We took the reward value on the immediate lever to depended linearly on the number of pellets received, *n*_*p*_, as *R*_*I*_ = *R*_3_ + (*n*_*p*_ − 3*Δ*R*, where *R*_3_ represents the perceived reward value for 3 pellets (the initial starting value) on the immediate lever relative to the constant reward on the delay lever, and Δ*R* the value of a single pellet. Note, that during the simulation, the agent receives the actual number of pellets currently assigned to each lever. We used a simple Q-learning update rule for the Q-value of the immediate lever, given by

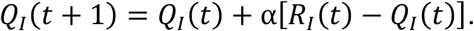

Given an initial lever bias, *p*_*D*_, the initial conditions are set so that *Q*_*I*_(0) = *R*_3_ = *log*((1 − *p*_*D*_)/*p*_*D*_*. Thus, in the absence of any resource tracking (i.e. Δ*R* = 0), or α = 0, the subject would just continue to pick levers with the initial bias set by *p*_*D*_. The subject then chooses levers according to the current Q-value of the immediate lever using the policy, *p*_*D*_(*t*) = 1/(1 + *e*^Q*I*(*t*)^*. Levers were chosen at random using the current value of *p*_*D*_(*t*) and Q-values and ival updated according to the task rules. We varied both Δ*R* and α at different initial lever bias, *p*_*D*_, and recorded the total number of pellets received.

### Analysis of theta oscillations in local field potentials

Local field potentials (LFPs) were acquired in each animal. For analysis, the 64 LFPs were averaged, and analysis was performed on this signal from each recording. Signals were down sampled to 1000Hz and the time around each choice (-10 sec: 20 sec) was extracted. Spectral decomposition was performed via short-time Fourier transform over 0.5 s windows with 90% overlap. Real components of the signal were extracted and power in the theta band was taken as the average power for each time bin in the 6-12Hz band. Power measures were smoothed via moving average over 50ms for each trial. To examine the impact of the value of the immediate lever on theta power, trials were stratified by low (0-2), medium (3-4), or high (5-6) pellets.

### Analysis of oscillations in neural firing

Autocorrelations in firing were computed for each neuron over +/-1s and binned at 1ms. PCA was then performed on autocorrelations (Figure 5A,B), and it was found that PC3 split the neurons that exhibited 4-5Hz or theta oscillations. This was validated by examining the spectrum (via Fourier transform) of the mean autocorrelation obtained from autocorrelations with either positive or negative coefficients. An examination of the distribution of coefficients associated with PC3 yield a distribution with three clear modes. Positive loading neurons (>0.015) exhibited 4-5Hz oscillations and negative loading neurons <-0.015) exhibited theta oscillations. In addition, an intermediate group (loadings > - 0.15 and < 0.015) with no clear oscillations was observed.

Spike-field coherence was assessed by computing the coherence in the spectrum of the spike trains and LFP for a 1 sec window surrounding each spike via Fast Fourier Transform using getCoherence.m^36^. Spike triggered averages in spike-field coherence were generated for each neuron in a session and then averaged across sessions.

### Analysis of value correlates in neural firing

The spike count for a given neuron in each 200 ms time bin, *i*, around a lever press, *t*, (we took -4 s to 13 s) was fit using a mixed effect model given by

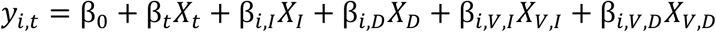

where β_0_ is the mean firing rate of the neuron, β_*t*_ accounts for any drift with trial number, *t*, with *X*_*t*_ = *t*, β_*i*,*I*_ and β_*i*,*D*_ characterize the lever press dependency with *X*_*I*_ = 1 on immediate lever trials, otherwise 0, and *X*_*D*_ = 1 on delay trials and 0 otherwise, and lastly, β_*i*,*V*,*I*_ and β_*i*,*V*,*D*_ characterize any additional ival dependence on either the immediate or delay levers with *X*_*V*,*I*_ or *X*_*V*,*D*_ being equal to the current ival on trial, *t*. If the t-statistic of β_*t*_ was significant (we used > 4, corresponding to a p-value ∼ 0.0001 and accounting for multiple testing of neurons), then the neuron’s firing was drifting over the session (potentially in a non-linear way) and we removed it from any further analysis. The set of fitted β’s for a neuron represent how its firing depends on lever press type and ival. We pooled together the β ‘s of all the neurons across all sessions in a given behavioral group. This pooled matrix was then clustered using hierarchical clustering to find the dominant models of firing across a group.

### Statistics

Where appropriate, groups effects were assessed via omnibus tests (Analysis of Variance, Kruskal-Wallace) followed by post-hoc tests with alpha = 0.05. Multiple comparisons corrections were applied via False Discovery Rate (Bejnamini and Hochberg’s method).

To assess the number of modes in a distribution, Gaussian Mixture Models were iteratively fit using fitgmdist.m (maximum iterations = 1000, tolerance function = 1×10^-6^) with an increasing number of modes from 1 to 5. Akaike Information Criterion (AIC) and Bayesian Information Criterion (BIC) were calculated for each mode. This process was repeated 100 times then values for AIC and BIC were averaged across bootstraps for each mode. Drops in AIC and BIC that reached an asymptote were taken as evidence for the optimal number of modes in the distribution.

To assess chance level of correlation between spike trains and ival, a bootstrapping procedure was implemented. This consisted of scrambling the sequence of ivals over trials for a session and correlating this with the intact firing rates. The process was repeated 1000 times creating a Pearsons Correlation Coefficient for each bootstrap that was used to calculate a 95% confidence interval to evaluate the upper bound of the randomized correlation. R values that exceeded this threshold were considered more than expected by chance.

## Results

The adjusting reward delay discounting (DD) task could theoretically be solved using some combination of two behavioral strategies that each requires the deployment of different forms of cognitive effort. The first strategy involves simply focusing on the delayed lever throughout the entire session. This is a non-impulsive strategy that would produce the largest overall pellet yield, but it requires a resistance-based form of cognitive effort to wait out the delays associated with each delay lever choice. If waiting out the delays are too aversive, the alternative strategy would be to shift between levers based on their relative value. This poses its own challenges as it requires a resource-based form of cognitive effort to continually monitor the number of pellets the immediate lever provides on each trial (Figure 1C).

**Figure 1:**
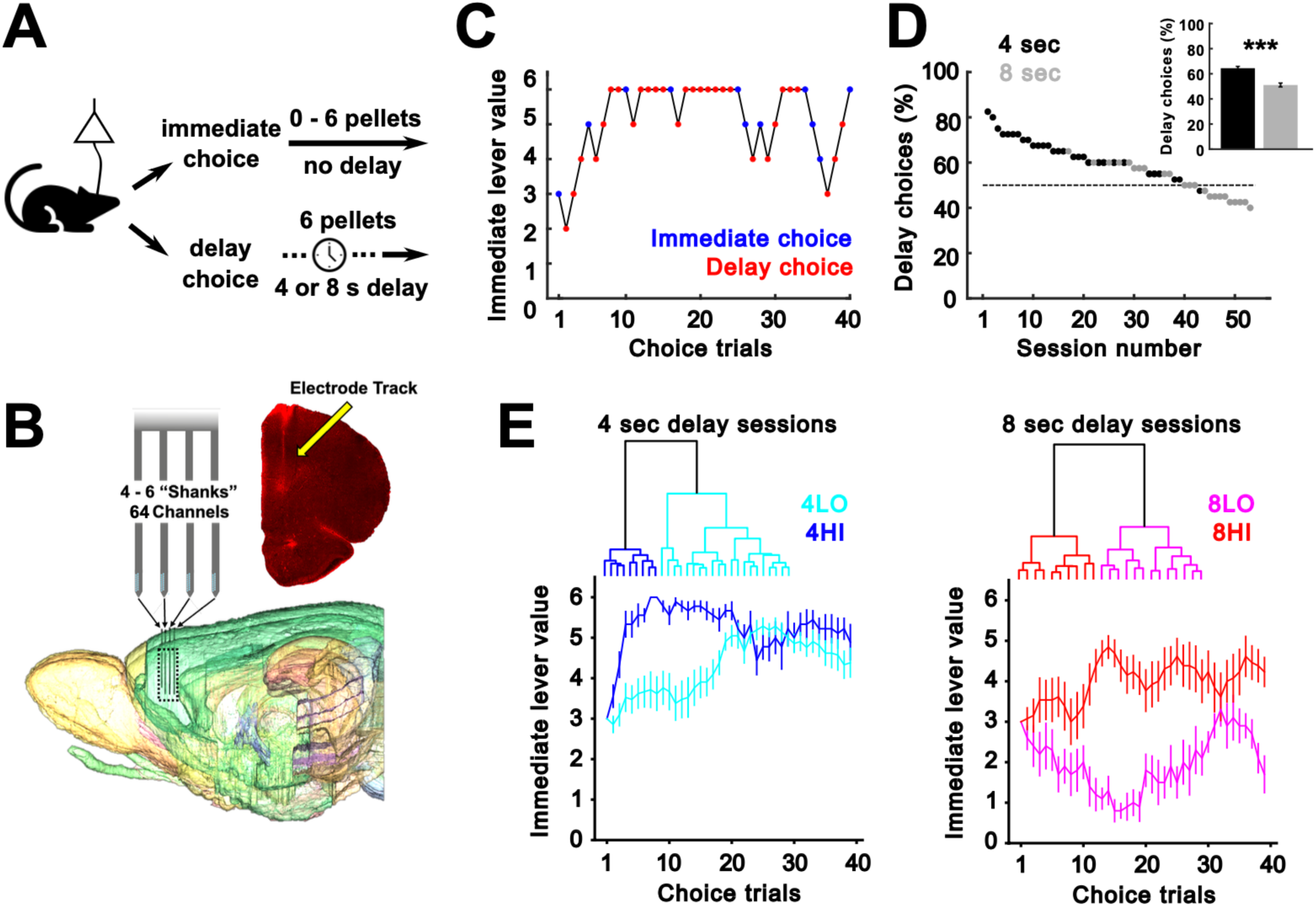
Capturing heterogeneity in choices across each delay. (A) Graphic describing the basic choice and outcomes in the adjusting amount delay discounting task. (B) Recording methods and example placement from a single animal. (C) Example session where the change in ival over trials is shown – decreases with immediate choices (blue) and increases with delay choices (red). (D) The percentage of delay choices (4s = blue, 8s = red) for each session in the data set. A decrease in the percent of delay choices is observed at the 8 sec delay (inset). (E) A hierarchical clustering model finds 4 distinct behavioral groups (top). The change in ival over the session is shown for each group (bottom). ***p<0.001.

**Table 1:** Description of session numbers from each rat in the behavioral groupings.

|  |  | Rat Number |  |  |  |  |  |  |  |
| --- | --- | --- | --- | --- | --- | --- | --- | --- | --- |
|  |  | 2 | 5 | 13 | 17 | 42 | 49 | 58 | 60 |
| Group | 4HI | 0 | 2 | 0 | 2 | 2 | 0 | 1 | 2 |
|  | 4LO | 1 | 1 | 1 | 3 | 5 | 3 | 5 | 2 |
|  | 8HI | 1 | 0 | 0 | 0 | 2 | 1 | 2 | 7 |
|  | 8LO | 0 | 0 | 0 | 0 | 0 | 7 | 3 | 0 |

### 4 behavioral groups are observed across all sessions

There was considerable heterogeneity in choice behavior at sessions employing either the 4s or 8s delay (Figure 1D). Across delays, choice reaction times tended to be fast (< 1 sec) but did not differ (Two sample t-test, t(2870)=1.32, p=0.19). While there was a clear shift towards choosing the immediate reward lever on sessions with an 8s delay [Figure 1D, independent samples t-test, t(52)=6.13, p=1.2×10^-7^], there were still sessions that delay lever presses were more prevalent. However, of the impulsive sessions, where the probability of choosing the immediate lever was > 0.5 (Figure 1D), all except 1, involved the 8 sec delay.

To capture this heterogeneity, we grouped sessions based on choice profiles at each delay. For this, hierarchical clustering on the observed patterns of free choice lever presses was performed. A level in the hierarchy was chosen so as to split sessions into 2 large groups at each delay interval (Figure 1E). The two resulting groups at each delay interval differed mainly in terms of how the value of immediate lever (ival) progressed throughout the session (Figure 1E). At the 4s delay interval, one group of sessions exhibited a consistent bias for the delay lever, which yielded a high sustained ival throughout the session (since presses on the delayed lever increased ival). This group of sessions will therefore be referred to as 4HI, for high ival sessions at a 4s delay interval. The second group of sessions started with a roughly 50/50 preference for the two levers which led to a relatively lower overall ival (Figure 1E). As a result, this group of sessions will be referred to as 4LO. However, note that the average ival of 4LO turned out to be above 3 because they focused more on the delayed lever later in these sessions (Figure 1E, bottom left).

At the 8 s delay, the two main clusters of sessions also differed in terms of ival, although the differences between high and low ival groups were starker. One group of sessions maintained ival above 3 (the 8HI group), while ival remained below 3 throughout the entire session in other group (the 8LO group). The 8LO group exhibited the strongest bias towards the immediate lever of any group at any delay (Figure 1E, bottom right).

### A delay bias is sufficient to capture choice distributions in the 4sec delay sessions

We then explored the factors that determined choice behavior in each group. A common feature of most groups was a bias for the delayed lever at some point in the sessions. Since most animals employed some mixture of strategies, it was difficult to quantify the impact of this factor in isolation. Therefore, we instead analyzed how differences in the strength of a delay bias impacted the choice behavior of an idealized simulated agent.

Even though there are only 2 lever options on this task, the total reward does not follow a binomial distribution because the payout on the immediate lever varies based on the choices made. Therefore, the simulations were set up such that the agent would choose stochastically but each choice was sampled using a constant bias parameter (see Methods). This parameter (*p*_*D*_), was based on the average probability of picking the delayed lever in each of the 4 groups described above. Figure 2A (left) gives the *p*_*D*_ histograms for each of the 4 groups, while Figure 2A (right) gives the resulting reward distributions over all free choices. Simulated agents choosing stochastically but with a *p*_*D*_ matching that of the 4HI group (*p*_*D*_ ∼ 0.73), produced a reward distribution that closely matched that observed for the 4HI group (KS-test, p-value = 0.72). This implied that the choice behavior of 4HI was exclusively dominated by *p*_*D*_ and appeared to benefit little by learning from prior outcomes. At the other extreme was the 8LO group, which had the lowest average probability for picking the delay lever (*p*_*D*_ ∼ 0.45). Simulations using this *p*_*D*_ produced a reward histogram that deviated from what was observed in the 8LO group (Figure 2A, KS-test p-value ∼ 0.1). In particular, the simulated agent would frequently achieve less reward than the 8LO group. This result means that the choice behavior of 8LO was not determined by a strong lever bias and suggested that this group must have relied on the alternate strategy of actively updating their choices based on prior outcomes (this will be explicitly tested below). Simulations involving the 4LO and 8HI were in between these two extremes.

**Figure 2:**
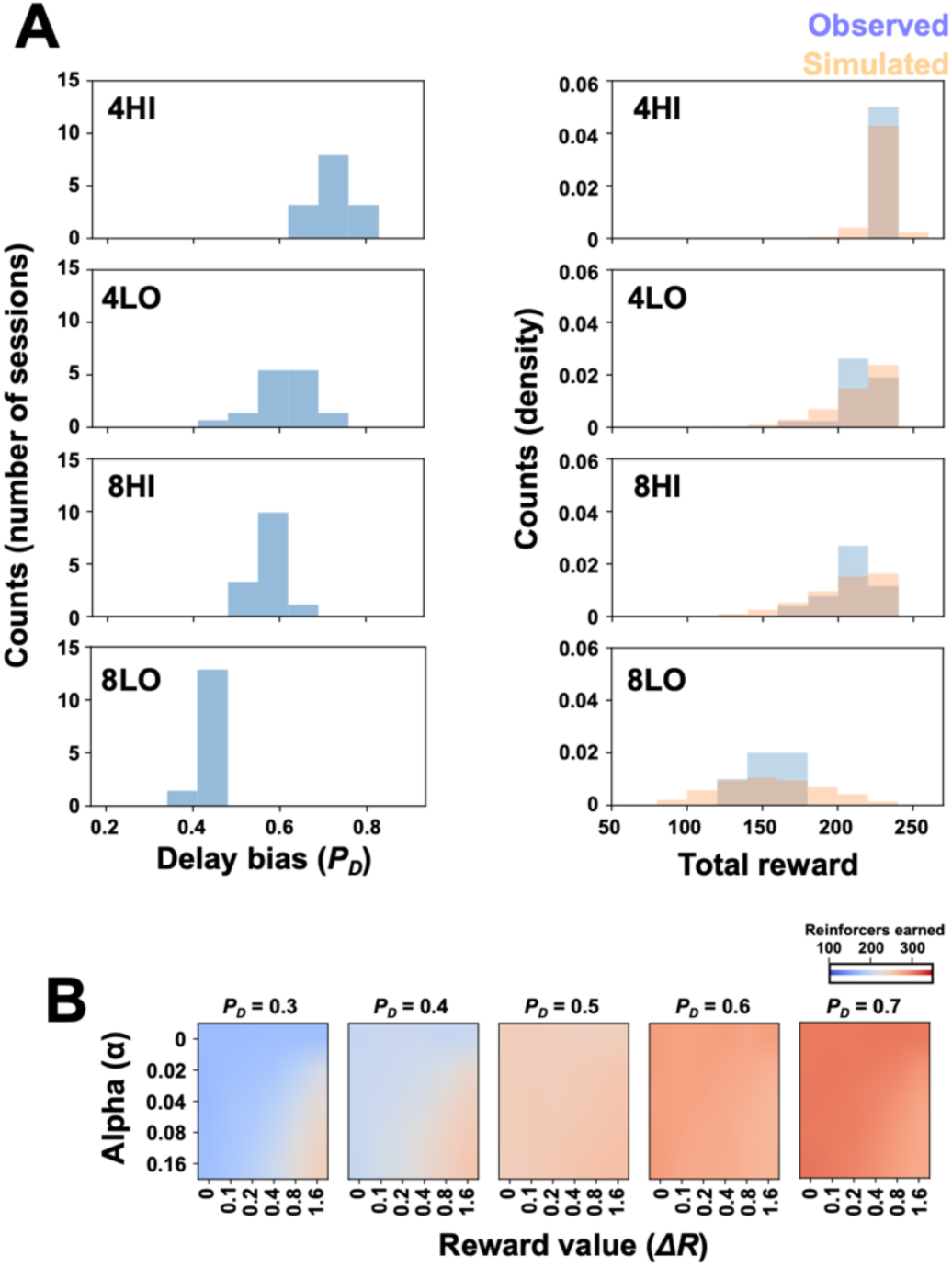
Simulations identify two main strategies that influence choice behavior. (A, left) The number of sessions in each cluster expressing a given delay bias (*P_D_*). (A, right) The total reward (pellets received) for the actual sessions (blue bars) and for the stochastic model simulations (light orange). For the simulations, *P_D_* was matched to the average *P_D_* of each of the real session clusters. The simulations produced similar pellet yields to those observed in the 4s session clusters (top) but a poor fit to the pellet yield in the 8 s session clusters (bottom). (B) A Reinforcement-Learning architecture was added to the basic stochastic model simulations and the relationship between ΔR (the change in the relative value of a single pellet with ival) versus Alpha (the rate at which the value of a pellet on the current choice affects subsequent choices) on the number of reinforcers (pellets) was plotted as a heatmap. A low ΔR means that the simulated agent valued a pellet the same regardless of ival, whereas the opposite is true when ΔR is high (i.e. ival-tracking). High PD produces the highest pellet yield overall, but pellet yield can be salvaged by a high ΔR when PD is low.

### Tracking value results in more rewards only when the delay bias low

As noted above, the alternate strategy to a delayed lever bias was to update choices based on prior outcomes, specifically ival. This strategy could be simulated by adding a simple reinforcement learning (RL) component to the model. Specifically, a Q-learning^37^ framework was employed where a Q-learning update rule adjusted the values of the choices of the simulated agent based on a valuation of the rewards received (see Methods).This approach follows from the work of others that have used Reinforcement Learning (RL) models to understand how the ACC may implement cognitive control^26,38–42^.

Our model now has three parameters: an initial bias towards the delay lever (*p*_*D*_), a learning rate (α), and a parameter that quantified the strength of resource tracking (Δ*R*). Figure 2B, shows the results of simulations using the updated model. In Figure 2B, the average total amount of pellets received on free trials is shown for different levels of *p*_*D*_ and Δ*R*. Overall, the total amount of reward increased with increasing bias, *p*_*D*_. However, note that at low *p*_*D*_ (< 0.4), the total reward increased with the strength of Δ*R*. This means that in the absence of a strong delayed lever bias, keeping track of ival would allow the animal to know when it is advantageous to switch to the immediate lever. This strategy cannot only yield reasonable reward payouts but is also valuable in the sense that it helps the animal avoid waiting out the delays imposed by the delay lever, particularly on sessions with an 8s delay interval. Conversely, at high *p*_*D*_ (> 0.5), there is minimal benefit from tracking ival, and in fact, the agent does slightly worse by using this strategy when Δ*R* is high.

Collectively the simulation results highlight the key factors that influence the two potential modes of cognitive control: a resistance-based form that is required to wait out long delays (formalized as a high *p*_*D*_), versus a resource-based form that involves tracking ival and switching to the immediate lever when ival is high (formalized as a high Δ*R*). By extension, the simulations make two strong predictions: Given that the behavior of 4HI could be replicated solely based on *p*_*D*_, there should be a clear neural representation of a resistance-based strategy in the 4HI group. Conversely, a resource-based strategy was optimal in simulations where *p*_*D*_ was low and impact of the resource tracking (Δ*R*) parameter was high. In this case, the choice behavior of the simulated agent placed a greater emphasis on immediate choices, similar to the 8LO group. Therefore, we should see a clear neural correlate of Δ*R* (i.e. ival tracking), in this group.

### Theta power increases on delay choices at the 4 sec delay

There is a growing interest in the role of frontomedial theta in cognitive control^26–29^. These studies typically involve macroscopic measures of cortical activity, such as EEG. This motivated us to explore whether neural activity patterns possibly reflecting an effort control signal could be observed in the spectrum of the local field potentials (LFPs) and spike trains within the rat ACC.

An assessment of the power spectrum in the LFP’s around the choice revealed that increases in theta that were most prominent on delayed lever choices at the 4 sec delay (Figure 3A). Further stratifying by the behavior groups revealed that theta power prior to the choice was most prevalent in the 4HI group (Figure 3B). Increases in theta power were observed in 4HI immediately prior to delay choices but were absent prior to immediate choices (Figure 3B1; Two-way ANOVA, choice x time interaction, F(590, 186756)=1.68, p=1.56×10^-22^). Following a 4 sec delay choice, theta power remained steady during the delay interval, dropped off at the termination of the delay, and then increased following the receipt of reward (Figure 3B1-2). Theta power prior to the delay choice was also transiently increased in the 4LO group (Figure 3B2; Two-way ANOVA, choice x time interaction, F(590, 443841)=6.73, p<1.0×10^-23^). The peak in LFP theta power would be consistent with what would be expected of a resistance-based form of cognitive control as it was present mainly in the group that focused on delayed lever presses (4HI) and it emerged prior to the choice when these rats were preparing for an impending delay.

**Figure 3:**
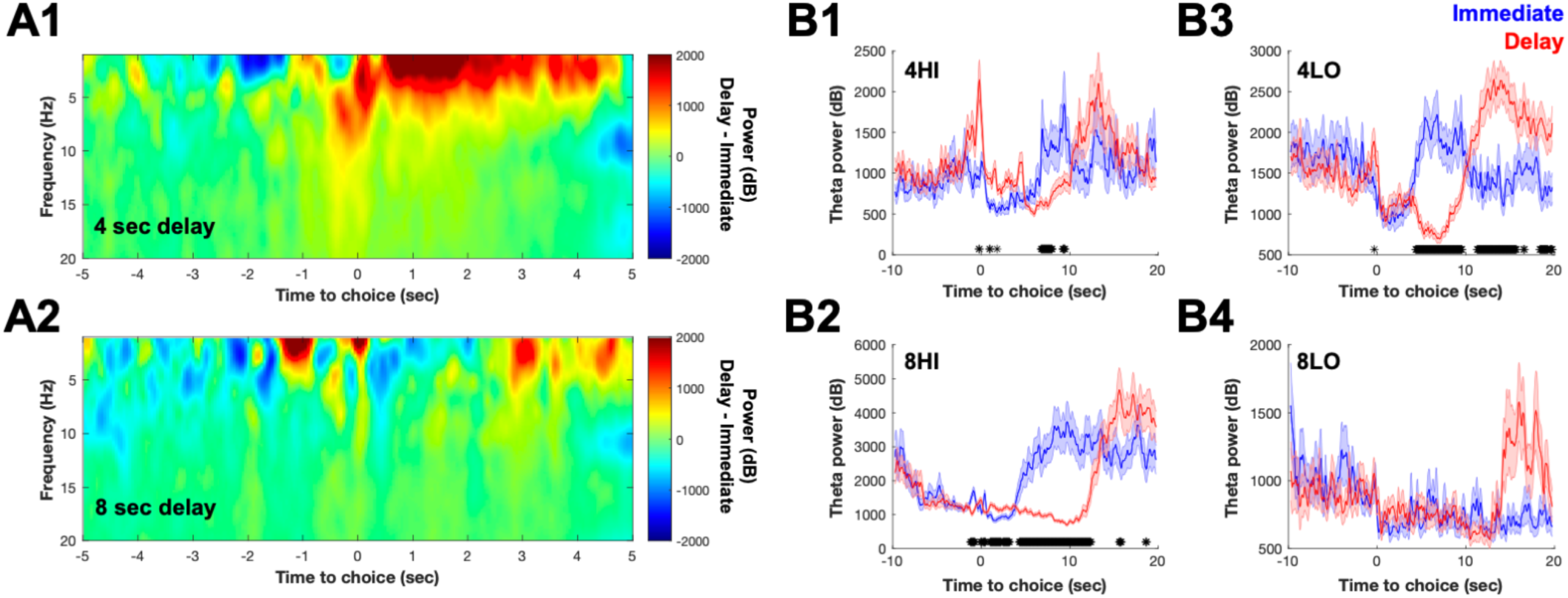
Theta power increases prior to 4s delay choices. (A) The difference spectrum up to 20Hz relative to the time of choice is shown for the average of the 4 s delay sessions (A1) and 8 s sessions (A2). Notice the increase in theta oscillations in proximity to the choice in the 4 s sessions (A1). (B) Theta power split by each of the behavioral groups and choice type. Robust increases in theta power are observed in the 4HI group immediately prior to the choice (B1), which is blunted in the 4LO group. Increases in theta power are not observed prior to a delay choice in the 8HI (B3) or 8LO (B4) groups. Time series are mean ± SEM. * p<0.05 FDR corrected rank sum tests.

If theta power in LFPs reflects a resistance-based signal, why would it be missing in the 8s groups where the delay intervals which were twice as long and hence placed higher demands on resistance-based control? The reason was because these groups largely avoided the delayed lever (Figure 1) and thereby abandoned the need to implement resistance-based control altogether. However, this in turn implies that the 8s groups should place a greater emphasis on the alternate, resourced-based control. Since this strategy involves tracking and using information about lever value (Figure 2), we first evaluated whether there was clear evidence of ival tracking in the 8s delay groups and second, whether such a signal was distinct from the theta-based signal observed in the LFPs (Figure 3).

### Neural correlates value tracking emerges prior to choice in the 8LO group

Similar to the changes in theta power, we examined neural activity around the choice for evidence of, in this case, ival tracking (see Methods for details). Briefly, for each neuron in every session, we fit the firing at each time bin around the lever press using a mixed effect model that depended on the lever press type (immediate or delay) and the current pellet value of the immediate lever (ival). The result of the fit was a model for the firing of each neuron around immediate or delay lever presses at varying levels of ival. The β values of each neuron in a behavioral group were pooled and then clustered to identify the dominant firing modes.

The prediction made above was that the strongest ival tracking signal should be found in the 8LO group. Figure 4A,B shows the cluster of neurons that provided the clearest example of value tracking in the 8LO group (permutation test for cluster compactness and separation, p-value < 1e-3). For each immediate choice, the averaged firing rate in a 2 s window around the lever press for neurons in this cluster was plotted against ival. Five example sessions are shown in Figure 4A. From these examples, clear correlations between neural activity and the value of the immediate lever are observable, consistent with predictions.

**Figure 4:**
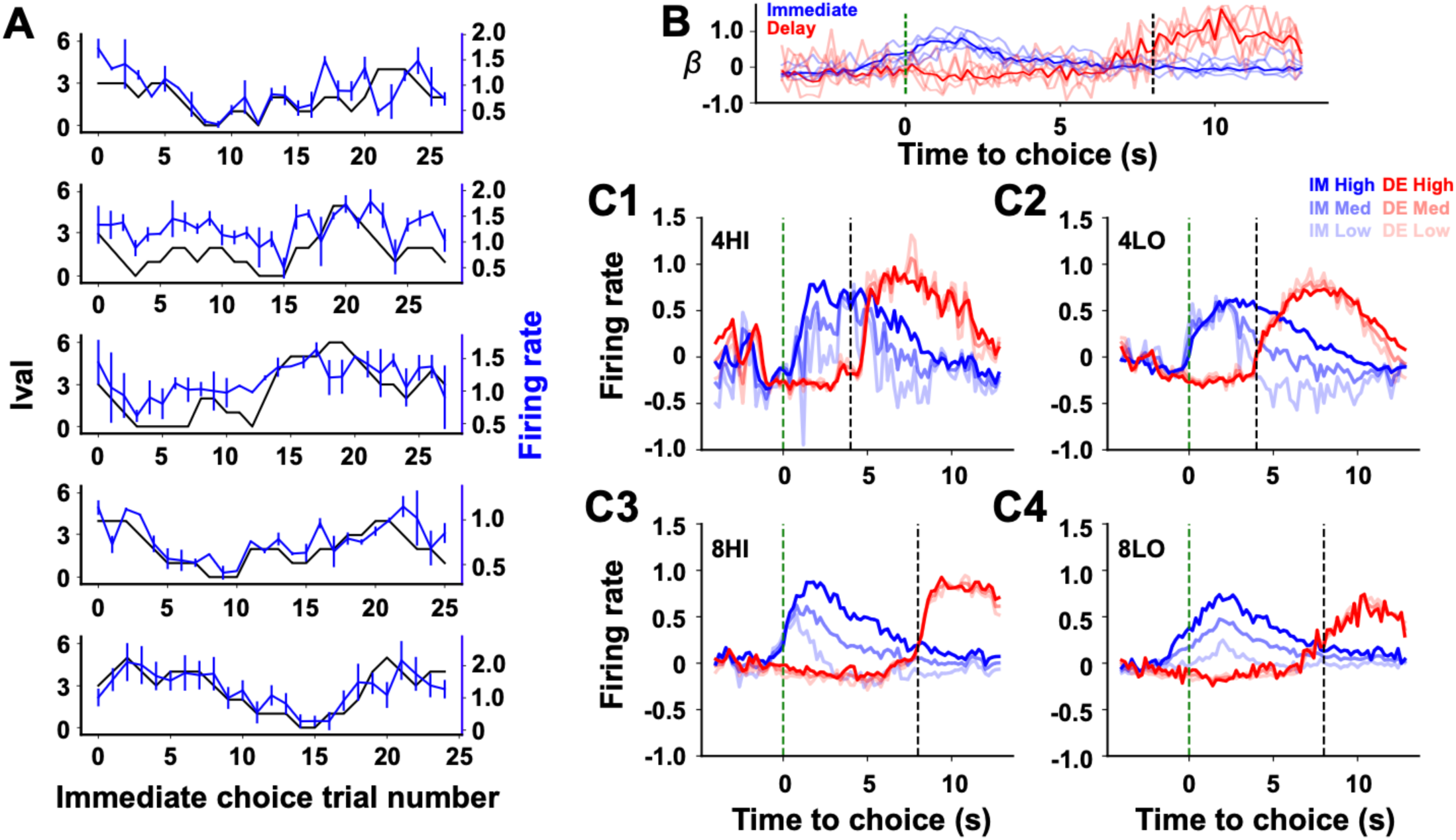
Ival-tracking neurons are prominent at 8 s delay. (A) The mean firing rate (blue lines) across all neurons in the ival tracking cluster is shown with the ival at each immediate choice (black lines) from five example sessions. (B) The mean beta values (β) at each time point from the neurons in the ival cluster (heavy lines). The β values for the ival cluster from each session in panel A are also shown (transparent lines). (C) The mean firing rate stratified by ival and choice type across the 4 behavioral groups. The dashed vertical green lines at time = 0 reflect the lever press time of the choice. The dashed vertical black lines denote the end of the delay.

Figure 4C stratifies the resource tracking cluster for each of the 4 behavioral groups and plots neural activity relative to the choice to determine *when* neurons in this cluster fire. On delay lever presses (blue curves), neural activity is independent of ival. However, on immediate lever presses (red curves), neural activity scales with ival. In addition, neural activity associated with ival depends on behavioral group; with 4 sec delay groups not having much variation in firing from low to high ival (Figure 4C1-2, whereas the 8 sec delay groups show much larger variation with ival (Figure 4C3-4). Indeed, in the 8LO group, neurons corresponding to the value tracking cluster were found in all sessions, whereas in the 4HI group, they came from only one of the nine sessions, suggesting this behavioral group had little to no resource tracking. Furthermore, the 8LO group exhibited ival-dependent increases in firing prior to the choice (Figure 4C4). Notably, this activity pattern was almost entirely absent in the other behavioral groups. The fact that ival encoding was observed prior to the choice in 8LO further indicated that value signals emerged at a timepoint when the actual lever decisions were being made.

### Theta coupled neurons do not track ival whereas non-theta coupled neurons do

Given the evidence for a neural signature of resistance (theta modulation) and resource (ival-tracking) based control, the final question was whether the two underlying processes were separate or interactive. To address this question, we first identified the neurons that were theta coupled by performing PCA on the spike train autocorrelations of each neuron regardless of behavioral group. PC3 was found to be the PC that most robustly separated out theta oscillating neurons across all sessions. Peaks in the power spectrum of autocorrelations were observed in the 4-5Hz and theta bands (Figure 5A-B). Neurons with positive loadings on PC3 tended to have autocorrelations whose power peaked at 4-5Hz (i.e. 4Hz coupled neurons), whereas neurons with negative loadings tended to have autocorrelations that peaked at theta (i.e. theta coupled neurons) (Figure 5B3,C). There was also a large swath of neurons with autocorrelations that lacked power at any frequency (No Oscillation neurons; Figure 5C). The existence of 3 distinct modes in the distribution of loadings was confirmed using Bayesian Information Criterion (BIC) and Akaike Information Criterion (AIC) (Figure 5C, inset).

**Figure 5:**
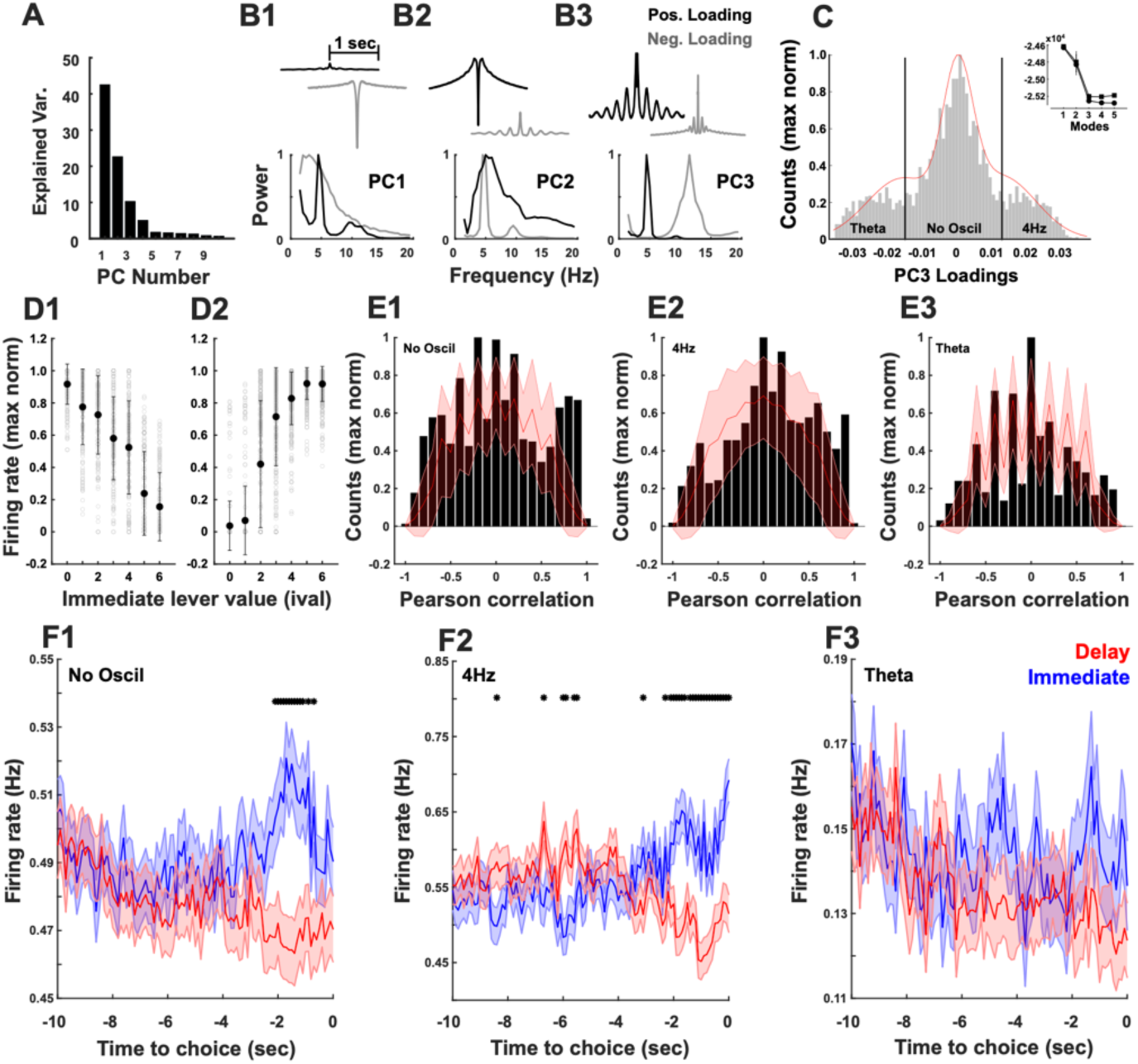
Theta and ival tracking neurons are distinct neural correlates. (A) The explained variance of each PC resulting from PCA performed on the spike train autocorrelations of all neurons in all sessions. (B) The raw spike train autocorrelation derived PC (top) and its power spectrum (bottom) captured by neurons with positive (black lines) and negative (gray lines) on each of the first 3 PC’s. Notice that PC3 identifies a well-separated pool of neurons whose autocorrelations have high power in the theta band (B3). (C) The distribution of neuronal loadings on PC3 exhibits 3 distinct modes, Theta-coupled, No Oscil, and 4Hz-coupled. The existence of three modes was validated by a drop in AIC and BIC at 3 modes that is asymptotic (inset). (D) The normalized firing rate versus ival for of neurons that exhibited a significant Pearson’s R for negative (D1) and positive (D2) correlations. Gray circles are from individual neurons, and circles are the mean firing rate ± SEM. (E) The count distributions of the number of neurons expressing a given correlation between firing rate and ival (black bars). The mean (red line) and 95% confidence interval (red shaded) of correlations calculated following shuffling ival over 1000 bootstraps. Distributions are shown for the No Oscillation group (E1), 4Hz Oscillation group (E2), and Theta oscillation group (E3). (F) The trial averaged firing rates leading up to the choice (0 sec) of the No-oscillation (F1), 4Hz (F2), and Theta (F3) -coupled groups. Time series are mean ± SEM. * p<0.05 FDR corrected rank sum tests.

If resistance (theta modulation) and resource (ival-tracking) based control were separate, the theta coupled neurons should not be influenced by changes in ival. Pearson’s correlation coefficients (R) between firing rate and ival was calculated for each neuron (examples in Figure 5D) and the distribution of R values in the theta coupled, 4 Hz coupled, and No Oscillation neurons are shown in Figure 5E. To evaluate if more R values than expected by chance approach +/- 1 in each group, we generated a 95% confidence interval (red lines, Figure 5E) based on shuffling ival and recalculating R (see methods). This analysis revealed that several neurons in the No Oscillation group exhibited extreme R values (Figure 5E1) that exceeded what was expected by chance. This implies that ival-tracking was over-represented in the No Oscillation group (Figure 5E1). Neurons in the 4Hz group also exhibited slightly more positive correlations than expected by chance (Figure 5E2). On the other hand, the few extreme R values in the theta oscillating neurons suggest that they are a distinct group that do not track ival (Figure 5E3).

The firing rate of neurons in the No Oscillation (Figure 5F1) and 4Hz (Figure 5F3) group differed in the way they responded on immediate versus delay choices, whereas neurons in the theta group did not (Figure 5F2; Two-way ANOVAs, choice x time interaction, No Oscillation, F(299, 1788600)=7.42, p= 0.19×10^-289^; Theta F(299, 4644000)=3.92, p= 4.99×10^-103^; 4Hz F(299, 403800)=39.75, p= 0). Collectively these data in Figure 5 indicate that it was the No Oscillation neurons that most robustly tracked ival and exhibited the strongest lever differentiation.

### Value tracking neurons are most prevalent in the 8LO group

If the No Oscillation neurons are ival-tracking then they should be enriched in the 8LO group that was most strongly biased towards a resource-based strategy. In the 8LO group, a larger proportion of neurons was indeed found to be of the No Oscillation, ival-tracking subtype (Figure 6A-B; No Oscil Group, Kruskal-Wallis Test, X^2(3,49)=8.28, p<0.04; Theta and 4Hz groups, Kruskal-Wallis Tests, p’s>0.5). In addition, a similar result is observed when the number of neurons (rather than proportion) in each group was calculated (Figure 6C). Conversely, 8LO had the fewest neurons with strong theta-oscillations, as predicted (Figure 6C). Each distribution was compared and each contrast with the 8LO group was significant (Figure 6C, X*^2^* Test of Independence, p’s < 0.1×10^-9^). In addition, when we measured spike-field coherence in each group and integrated power in the theta band for each neuron (Figure 6D1-2), theta spike field coherence was again the lowest in the 8LO group (One-way ANOVA, F(3,4110)=37.33, p=8.58×10^-24^). These data revealed that ival tracking was most prominent in the 8LO group.

**Figure 6:**
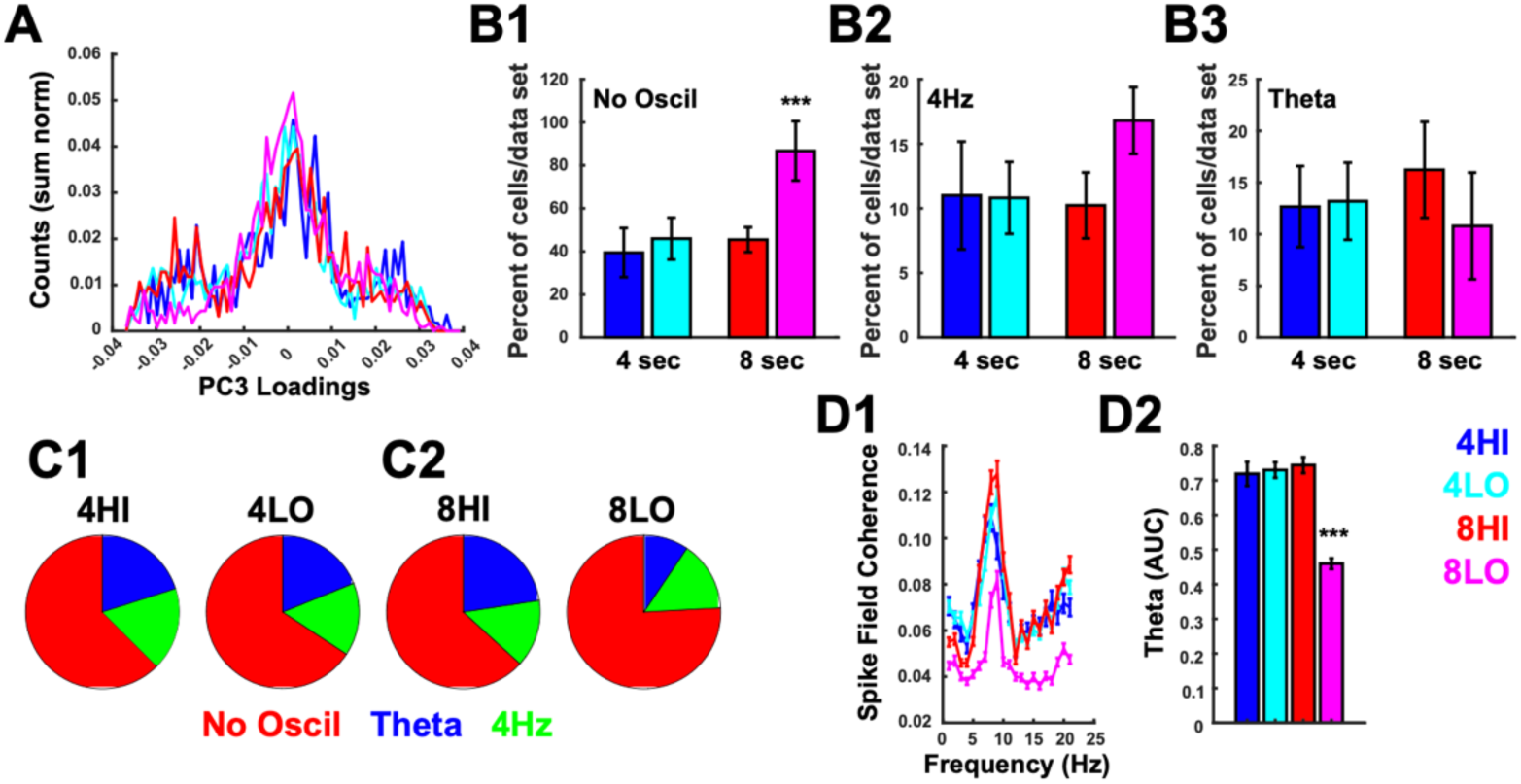
The behavioral groups exhibit distinct neural correlates of effort strategies. (A) The neuronal loadings of the autocorrelations on PC3 stratified by the behavioral groups (4HI=blue, 4LO=teal, 8HI=red, 8LO=magenta). The percent of neurons No-Oscillation neurons (B1), 4Hz-coupled neurons (B2) and Theta-coupled neurons (B3) stratified by behavioral group (4HI=blue, 4LO=teal, 8HI=red, 8LO=magenta). (C) The total number of No-Oscillation neurons (red), 4Hz-coupled neurons (blue) and Theta-coupled (red) neurons stratified by behavioral group. (D1) Spike-field coherence up to 20Hz is shown for each of the behavioral groups (D1). (D2) The area under the curve for the theta band is stratified by the 4 behavioral groups. All data are mean ± SEM, *** p < 0.001, post-hoc tests.

Collectively these data indicate that theta oscillations, and the role they might play in resistance-based effort, are blunted in the 8LO group, whereas ival-tracking is enhanced. This is in stark contrast to the 4HI group that exhibited the strongest bias towards the delay lever, the largest portion of theta-tracking neurons and the fewest ival-tracking neurons, as described above. These data indicate that there are two separate strategies at play on this task that are mediated by two largely independent neural mechanisms. Prominent pre-choice theta power was associated with a behavioral strategy characterized by a strong bias towards a resistance-based strategy, whereas the neural signature of ival-tracking was associated with a strong bias towards a resource-based strategy.

## Discussion

In the present study, the adjusting amount delayed-discounting task was used to search for neural signals in ACC related to cognitive control. Sessions were clustered into 4 main groups, based on delay duration and the propensity to make immediate choices. A RL-model framework was used to reproduce the choice patterns of each group, which indicated that either a resistance or resource - based effort control strategy provided a sufficient description of the behavior on this task. A neural signal related to ival-tracking was clearly observed, providing a candidate for a resource tracking signal. In addition, a putative resistance-based signal could also be found at the single neuron and LFP level but only short delays where delay choices were highly prevalent. These results provide insights into how ACC networks might encode cognitive effort in the service of cognitive control.

One way to define cognitive effort is in terms of the domain that is being taxed (e.g. attention, memory, problem solving). However, it can also be defined in a more descriptive sense, emphasizing the process rather than the domain. In this framework, a *resourced-based* form of cognitive effort is utilized whenever a valuable but limited capacity resource, such as attention or working memory, is used to solve a task^1,3,4^. In contrast, a *resistance-based* form of cognitive effort is used in order to overcome some type of internal ‘resistive force’, such as unpleasantness or impatience^2,5^.

Typically, a resistance-based form is most relevant to delay-discounting tasks as effort is needed in order to wait out delays and resist the temptation of the immediate reward^43,44^. However, both types of cognitive effort are required for the adjusted amount delay-discounting task because this task incorporates the dynamic variable ival, which fluctuates throughout the session based on the animal’s behavior. A food-deprived rat seeks to both maximize reward and avoid delays. A resistance-based form of cognitive effort is needed if rats opt for a dLP-biased strategy because this forces them to wait out all the delays associated with the delayed lever. Alternatively, the rat could keep track of ival in order to know when the payout of the immediate lever was high enough to warrant a switch from the delayed lever and thereby avoid the associated delays without significantly sacrificing pellet yield. However, ival-tracking strongly taps a resourced-based form of cognitive effort since it places a high demand on attention and mnemonic resources. Here we found that the resistance-biased strategy predominated when the delays were short (4s), whereas a resource-based strategy predominated when the delays were long (8s).

The model results suggested that a simulated RL agent could solve this task by simply focusing on the delay lever. However, this was not sufficient to accurately fit the distribution of choices in all behavioral groups, which in turn implied that an additional strategy was at play. This strategy involved tracking ival and was formalized by the parameter Δ*R* in the model. Consistent with model predictions, we found a strong neural signal within the ACC that closely tracked ival and was most prevalent in the 8 s delay sessions. Further, this signal encoded the value of the immediate lever prior to the choice in the 8LO group. This is consistent with the conclusion that this group deployed a resource-based form of effort to guide lever choices on this task.

The present results provide a mechanistic basis for theories highlighting a role of the ACC in cognitive control^1,10,45,46^. Cognitive control processes, mediated by the ACC, are thought to dynamically regulate how much and what type of cognitive effort should be deployed based on the expected value of the outcome, relative to the cost of implementing and maintaining the cognitive effort. Indirect support for this theory comes from human imaging studies^47^ and electrophysiology studies in primates^48^ and rodents^49^ showing that ACC neurons encode the value of outcomes. ACC neurons potentially compute value by combining information about the magnitude of a reward^50^ and the spatial or temporal distance to it^51,52^. There is also clear evidence for a role of the ACC in signaling effort. Lesions of the ACC make it less likely that rodents will choose an option that requires greater physical effort, even if that effort yields a higher reward ^20–22,53^. Inactivation of ACC in the rodent was also found to reduce the willingness of rats to expend cognitive effort as defined in terms of visuospatial attention on a variant of the 5-choice serial reaction time task^23^. In addition, neurons within the rodent and primate ACC signal the degree of physical effort, regardless of whether effort is defined in terms of the size of an obstacle, the angle of a ramp to be traversed, or in terms of competitive effort^17,30,48,54,55^. Most of these studies have found that ACC neurons encoded a multiplexed representation that combined information about relative effort and the relative value of the reward. Hillman & Bilkey^56^ referred to this multiplexed signal as representation of the overall net utility, which fits well with the description of value in the theories of cognitive control mentioned above.

Holroyd and colleagues^25,28^ investigated potential electrophysiological correlates of cognitive effort in human EEG signals. They built on findings suggesting that the ACC is a main generator of ‘frontal midline theta’ and argued that this signal is related to the application of both physical and cognitive effort. They postulated that when this signal is combined with a second reward-related signal (also believed to be generated within the ACC), it creates a representation of expected value or net utility. Their theory is supported by fMRI studies that have found that representations of anticipated effort and prospective reward elicit overlapping patterns of activation in ACC^19,57^. Our findings in rodent ACC fit nicely within this framework. We speculated that variations in theta characteristics could underlie a resistance-based form of control. Specifically, the fact that theta power was most robust in 4HI group and only observed prior to delay choices, fit the bill for a resistance-based control signal. Conversely, theta entrainment of neural activity was lowest in animals that used a resource-based control strategy (8LO).

Cognitive control theories are by definition, cognitive in nature. However, it is difficult to know whether ACC neurons compute value or cost in a cognitive/economic sense or whether the ACC simply tracks the internal reactions to the results of such computations performed elsewhere in the brain. On one hand, the ACC is part of a cognitive control network that includes “cognitive” regions such as the posterior parietal cortex and prefrontal cortex^58^. On the other hand, it has extensive, often bi-directional connections, with subcortical, brain stem and spinal cord regions involved in tracking and modulating emotional state and autonomic tone^14–16^. Accordingly, the main effects of ACC stimulation are changes in autonomic markers such as heart rate, blood pressure and breathing^16,59^. The integration of ACC signals in this array of brain regions would likely require a mechanism to synchronize neural activity amongst them and theta oscillations would be a good candidate. This may facilitate the integration of cognitive, emotional, and autonomic signal across brain-wide circuits.

However, the main problem in determining whether recorded signals are cognitive or affective in nature is that laboratory tasks, including the adjusted-reward delay-discounting task, use biologically relevant events as both a source of information and motivation. For instance, consider the observation that ACC ensembles robustly tracked ival. By tracking this variable, rats could in theory create a relative value representation of both levers, informing them about when the expected payout of the immediate lever was high and therefore when it was advantageous to switch to this lever. However, ival is also a proxy of the recent reward history. Therefore, the ival-tracking signal could be used to guide responses or could simply reflect the emotional or autonomic response to the ongoing tally of relative wins and losses. The same could be said of theta band synchrony. Prior work has implicated theta band synchrony in frontal cortex with either the amygdala or the ventral hippocampus during periods of stress and/or anxiety^60,61^. Further, a reduction in synchrony in theta between these regions corresponds to a loss of control over stress and anxiety^60,62,63^. While it is difficult to equate 4 or 8 sec delays with an anxiety or stress producing stimulus, it is clear that the animals do not “like” the delay as they will avoid the delay option whenever possible. Theta synchrony may generate a resistance-based control signal that is deployed to mitigate the unpleasantness of the delay in this task.

Cognitive effort is costly because it is inherently aversive^4^. It has been argued that the cost of cognitive effort is ‘felt’ at a physiological level through changes in emotion and autonomic tone^2,4,12,13^. Therefore, ACC may assess the cost of cognitive effort in the same way it calculates the value of an expected outcome, which is indirectly via changes in autonomic tone. We would argue that this is generally the case, in that the primarily function of the ACC is likely to monitor and regulate autonomic tone^16^, but when these signals are transmitted to downstream regions, perhaps via theta oscillations, they serve as important cues that guide cognitive control and decision-making.

## ACKNOWLEGMENTS

The work was supported by grants from NIH (AA029409, P60-AA007611, and T32AA007462), CIHR (MOP-137045 and PJT-159796) and NSERC (05979).

## AUTHOR CONTRIBUTIONS

D.L., M.M., S.W., C.C., and C.L. designed the *in vivo* recording study. S.W., M.M., B.M. and D.L. conducted the *in vivo* recordings. J.S., EE, and C.L. conduced the data analyses. J.S., EE, and C.L. wrote the paper.

## COMPETING FINANCIAL INTERESTS

The authors declare no competing financial interests.

## Notes

### Competing Interest Statement

The authors have declared no competing interest.

### Summary of Updates

We are extremely grateful for the several excellent comments of the reviewers. To address these comments, we have completely reworked the manuscript adding more rigorous approaches in each phase of the analysis and computational model. Here is a (nonexhaustive) overview of the major changes we made: (1)We have developed a way to more adequately capture the heterogeneity in the behavior. (2) We have completely reworked the RL model. (3) We have added additional approaches and rigor to the analysis of the value-tracking signal.

